# Helminth infection favors reprogramming and proliferation of lung neutrophils

**DOI:** 10.1101/2025.03.25.645229

**Authors:** Fei Chen, Suheyla Batirbek, Vanessa Espinosa, Lianhua Jin, Keyi Wang, Wenhui Wu, Evan Johnson, Alexander Lemenze, Adriana Messyasz, Mark Siracusa, Dane Parker, Amariliz Rivera, William C. Gause

## Abstract

Neutrophils are a granulocytic population of myeloid cells that have critical effector functions during infectious disease but are generally thought to be short-lived and nonproliferative with markedly limited activation states. In these studies, we directly compared lung neutrophil activation following infection with different groups of pathogens including bacteria, fungi, and helminths. Our results demonstrate considerable heterogeneity depending on the type of infectious agent. In contrast to bacterial and fungal infection, after helminth infection neutrophils expressed markers associated with characteristic type 2 responses and unexpectedly upregulated genes associated with cell cycling and protein synthesis. Further studies showed reduced neutrophil cell death following helminth infection and increased proliferation, which was dependent on IL-4R signaling. This distinct subset of proliferating neutrophils expanded following helminth infection and was released from the endothelial niche to colocalize with invading parasites in the airways. These studies demonstrate a novel long-lived cycling phenotype for neutrophils following helminth infection.

---

Neutrophils are recognized as rapid-response effector cells of the immune system that can quickly attack invading pathogens with a diverse array of effector functions including phagocytosis and production of reactive oxygen species (ROS), release of antimicrobial proteins and DNA-based neutrophil encapsulating traps. However, unlike other immune cells, they are also generally thought to be short-lived and nonproliferative in extramedullary environments, with relatively low levels of RNA reflective of their specialized and short-lived activities^1–3^.

Following their release from the bone marrow into the blood, neutrophils have been estimated to persist in the circulation for less than a day^4^. Recent studies however indicate that neutrophils migrating to specific tissues can also rapidly develop a tissue-specific signature revealing a previously unrecognized plasticity in neutrophil activation that may be important in maintaining homeostasis, though still persisting for little more than a day^2^. However, in the context of the nonphysiologic tumor microenvironment (tme) recent studies suggest that neutrophils can proliferate and survive for at least several days^5, 6^. Most studies examining neutrophil effector functions during infectious disease have been performed in the context of prokaryotic microbial infections. However, recently neutrophil activation and effector function have been shown to also occur during fungal^7^ and helminth infections^8, 9^, though as yet few studies have examined whether their activation state differs relative to responses to prokaryotic microbes.

## Lung neutrophil activation varies in response to infection with different types of respiratory pathogens

We examined the possibility of plasticity in lung neutrophil activation associated with neutrophil lung infiltration that occurs after infection with distinct groups of pulmonary infectious pathogens. BALB/c mice, housed in the same animal facility to minimize variation in housing conditions, including microbiota, were infected with either helminths (*Nippostrongylus brasiliensis*), fungi (*Aspergillus fumigatus*), or bacteria (*Staphylococcus aureus*) in the same experiment, a timepoint at which neutrophils are markedly increased in the lung in response to infection with all three pathogens^10, 11^. Two days after infection, sort-purified CD45^+^, CD11b^+^, Ly6G^+^ lung neutrophils were subjected to global transcriptome profiling. It should be noted that we did not detect CD64^+^, Ly6G^+^ macrophages at this early time point (data not shown), consistent with this population arising at later stages of the response after Influenza infection^12^. Neutrophils isolated from each pathogen demonstrated a distinct profile of upregulated genes (relative to neutrophils from untreated lungs) in each pathogen group, as shown by principal component analyses (PCA) (Figure 1a). Gene expression patterns showed both unique and overlapping upregulated expression of genes across fungi, bacterial, and helminth infections (Figure 1b). These findings indicate the plasticity of the neutrophil transcriptome after acute infection that may enable neutrophils to rapidly develop tailored responses to these distinct pathogens. Using Ingenuity Pathway Analyses (IPA), predicted cell cycling signaling pathways were downregulated after SA infection while they were markedly upregulated following Nb inoculation, and more modestly increased after AF infection (Fig. 1c), with changes in specific genes shown in Fig. 1d. Predicted pathways associated with protein synthesis (eukaryotic translation initiation and elongation) were also preferentially activated in lung neutrophils from Nb inoculated groups (Fig. 1c). Immune response genes were also differentially upregulated in many cases mirroring responses seen in macrophages following infection with these different pathogens. For example *Chil3*, *Arg1*, and *Retnla* were elevated in neutrophils isolated in lungs after Nb infection, which are characteristic of alternative (M2) macrophage activation^13^, and the type 2 cytokine, IL-13, was also elevated. In contrast, neutrophils from SA infected mice upregulated genes more associated with type 1 immunity, including IL-12 and nitric oxide synthase 2 (NOS2), while neutrophils from AF infected mice showed a more mixed type 1/2 pattern. IPA showed similar predicted patterns of differential activation of type 1 and type 2 signaling pathways (suppl Fig. 1a). Remarkably, following Nb infection genes associated with xenobiotic pathways were selectively elevated (Fig 1d and suppl Fig. 1a), consistent with studies suggesting that type 2 immune responses may mediate protective responses to toxic or noxious stimuli^14–16^. Seahorse real-time cell metabolic analyses using the mitochondria stress test revealed a higher oxygen consumption rate (OCR) and a higher extracellular acidification rate (ECAR), in lung neutrophils from mice infected with Nb and AF, indicating overall increased metabolic activity and oxidative phosphorylation relative to neutrophils from SA infected mice (sFig.1b). Interestingly, increased mitochondrial activity has previously been associated with more immature neutrophils in the bone marrow as well as in the extramedullary tme^17^.

**Fig. 1.**
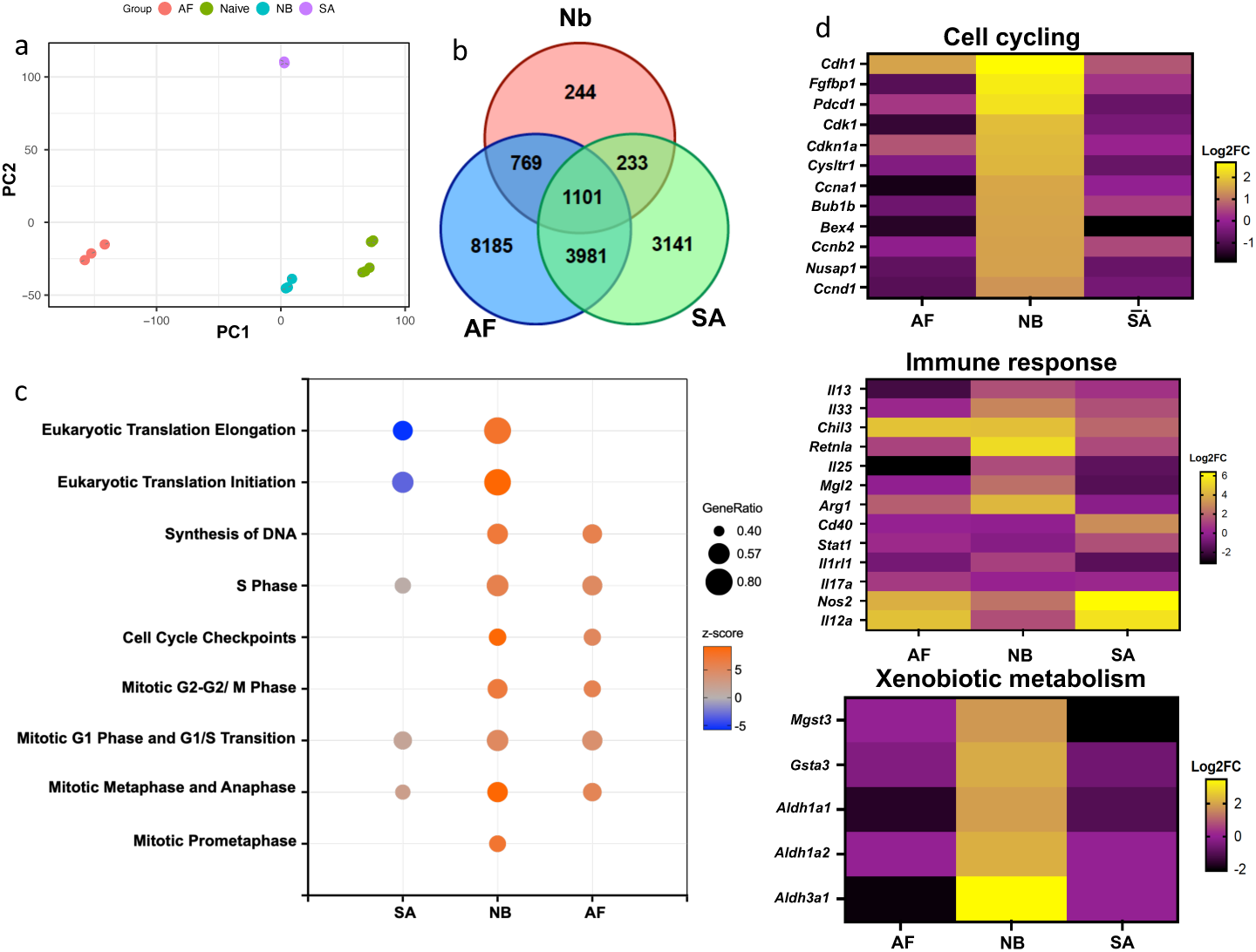
Lung neutrophil activation varies in response to infection with different types of respiratory pathogens Mice were inoculated with helminths, *N. brasiliensis* (Nb), fungi, *A. fumig*atus (Af), or bacteria, *S. aureus* (SA) for 2 days. Lung neutrophils were sort-purified for RNA-seq transcriptional analysis and compared to naïve neutrophils, with 3 mice/treatment group. (a) Principal component analysis (PCA) of the normalized RNAseq data transcripts. (b) Venn diagram illustrating overlap of upregulated or downregulated significant genes (FDR adjusted p-value <0.05) relative to neutrophils from naïve mice. (c) Bubble plot of Ingenuity Pathway Analysis (IPA) of differentially expressed genes (DEGs). “GeneRatio” indicates the ratio of enriched DEGs to total number of genes within the pathway. Z- score indicates activation status of the pathway. (d) Heatmap of selected characteristic markers related to cell cycling, immune responses, and xenobiotic metabolism in neutrophil responses to different pathogens, Nb, AF, and SA. Gene expression is shown as relative to neutrophils from naïve mice.

Increased oxidative phosphorylation is also characteristic of M2 macrophages, which also develop in the context of type 2 immunity and helminth infections^18–20^.

Taken together, these findings indicate the plasticity of neutrophils, specifically demonstrating their capacity to develop characteristic activation states tailored to distinct types of infectious agents invading pulmonary tissues. Surprisingly, lung neutrophils from Nb infected mice expressed genes associated with proliferation, raising the possibility that in the context of type 2 immunity neutrophils may preferentially undergo extramedullary cell cycling in the lung tissue microenvironment.

## Neutrophils from helminth-infected mice exhibit reduced cell death and increased cell cycling

Our findings from our transcriptome analyses indicated that neutrophils from helminth-infected mice preferentially expressed markers associated with type 2 immunity, wound healing, and also proliferation. Previous studies have shown that macrophages in the context of helminth infection express an alternatively activated (M2) macrophage state that includes activation of wound healing effector mechanisms^13^. More recently, in contrast to macrophages activated during type 1 responses these M2 macrophages have been shown to be capable of proliferating in extramedullary peripheral tissues during the helminth-induced type 2 immune response^21^. Neutrophils in the circulation typically have a half-life of about 12 hours^22^. Recent studies suggest that neutrophils in extramedullary tissue microenvironments can persist for longer periods, as long as several days^23^. Indeed, under physiologic conditions, and following infection, the non-proliferative state of neutrophils outside the bone marrow is considered a characteristic feature of this myeloid population. However, studies of neutrophils have been primarily conducted in peripheral blood or in the tissue microenvironment in either naïve mice or mice infected with microbial pathogens, including bacteria and viruses^2, 23, 24^. Given our unexpected findings that during helminth infection neutrophils showed elevated expression of genes associated with cell cycling, we next examined whether neutrophils activated in the context of type 2 immune responses and helminth infection showed reduced cell death. Lung cell suspensions were stained with Annexin V and propidium iodide (PI) at day 3 after infection with Nb, AF, and SA and analyzed by flow cytometry to identify early apoptotic (Annexin V+, PI-), early necrotic (Annexin V-, PI+), and late apoptotic/necrotic populations (Annexin V+, PI+), as previously described^25^. As shown in Suppl Fig. 2a, apoptotic and necrotic populations were markedly increased after SA and AF infection; however, elevations in apoptotic or necrotic neutrophils relative to naïve mice were not readily detected after Nb infection. To directly examine cell cycle progression after infection with these pathogens, at day 3 after Nb inoculation, mice were administered EdU i.p. 3 hours prior to sacrifice to assess cell cycle progression into S phase, as previously described^26^. As shown in Suppl Fig. 2b, a reciprocal pattern was detected compared to that observed with death pathways in Suppl Fig. 2a. Lung neutrophils from Nb inoculated mice showed significantly increased numbers of proliferating cells relative to lung neutrophils from naïve or either AF or SA infected mice. The increased number of cycling neutrophils observed in the lung after Nb inoculation may actually have resulted from neutrophils proliferating in the blood or bone marrow prior to their migration into the lung tissue. To examine this possibility, 3 days after Nb inoculation and 3 hours after EdU i.p. injection, lungs, blood, and bone marrow were collected and cell suspensions were prepared and stained for neutrophils (Ly6G) and ki67, a proliferation marker expressed at all stages of the cell cycle^21^. As shown in Suppl Fig. 2c, significant increases in EdU incorporation and Ki67 staining were detected in neutrophils in the lung but not in either the blood or the bone marrow.

## Neutrophils exhibit a long-lived proliferative phenotype after helminth infection

Nb exits the lung and enters the small intestine by day 4 after inoculation^27^. However, inflammation persists and can lead to emphysematous pathology by 30 days after infection^28^. To directly examine whether proliferating neutrophils persist in the lung after Nb inoculation, lung neutrophils were isolated by sort purification at day 2 after Nb inoculation of CD45.2 donor Balb/c mice and transferred intratracheally to infection-matched congenic CD45.1 recipient mice. Ten days later recipient mice were administered EdU i.p. and at day 11 blood and lung cell suspensions were assessed for neutrophil expression of EdU and Ki-67. As shown in Fig. 2a, although CD45.2 donor neutrophils were not detected in the blood, about 4% of lung neutrophils expressed CD45.2. Furthermore, these neutrophils stained positive for both EdU and Ki67, indicating that the surviving transferred neutrophils continued proliferating in the lung tissue microenvironment as late as 10 days after helminth infection. These studies demonstrate that neutrophils can persist and proliferate in lung tissue remodeled in the context of chronic type 2 inflammation for as long as 10 days. They also corroborate our findings in Fig. 2 that during a type 2 immune response neutrophils self-renew in lung extramedullary tissue rather than the bone marrow or blood tissues. Our studies now indicate for the first time that neutrophil proliferation occurs in the lung tissue microenvironment after helminth infection.

**Fig. 2.**
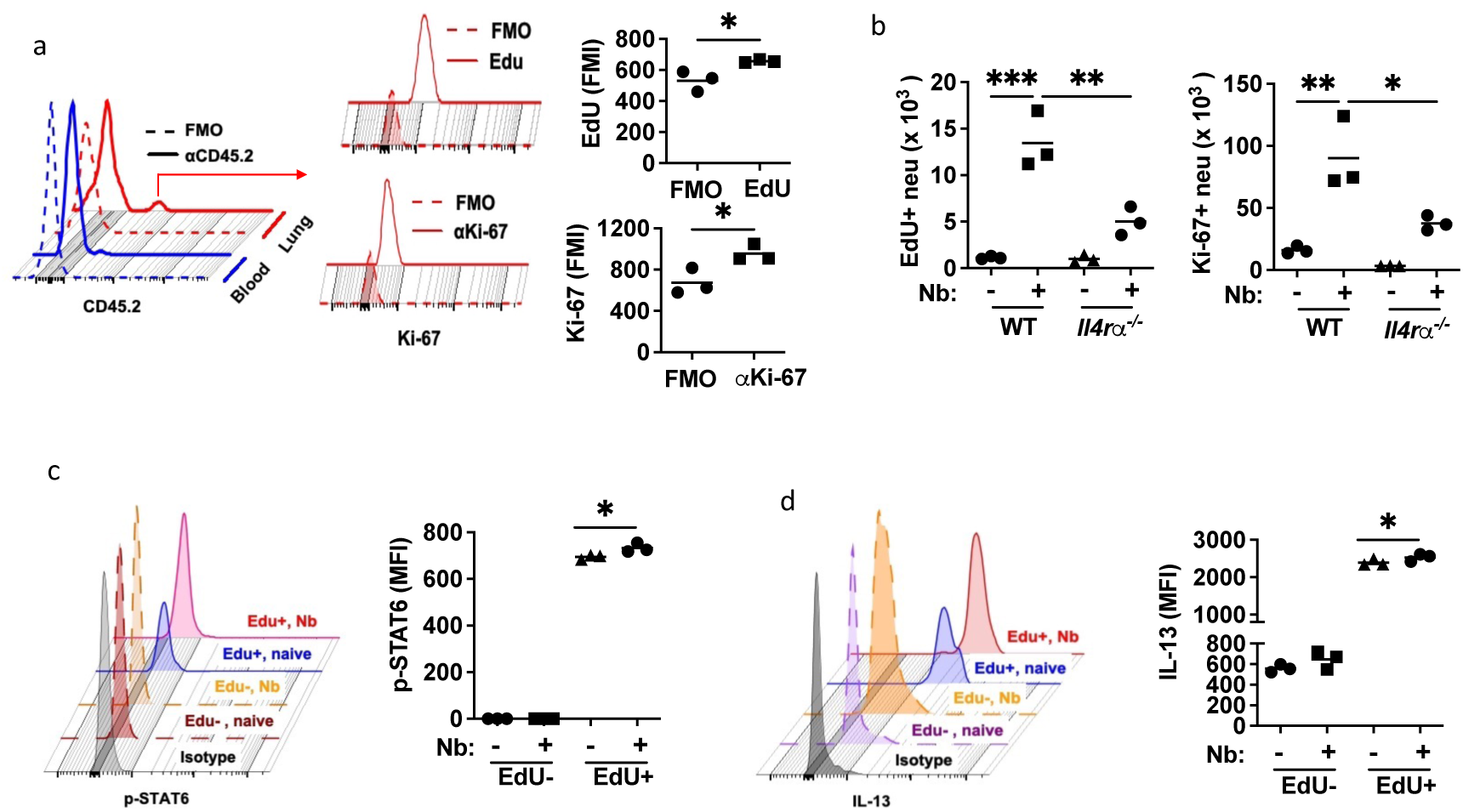
Neutrophils exhibit a long-lived proliferative phenotype after helminth infection, and IL-Rα dependent proliferation, IL-13, and p-STAT6 production (a) Infection-matched donor CD45.2 mice and CD45.1 recipients were inoculated with Nb for 2 days. Neutrophils (Zombieuv^-^CD45^+^CD11b^hi^Ly6G^hi^) from the lungs of donor mice were electronically sort purified and transferred i.t. into recipient mice. Ten days after cell transfer, recipients received EdU (i.p.) for 24 hrs, and donor gated CD45.2^+^ neutrophils from blood and lung tissue (gated on Zombieuv^-^CD45.2^+^CD11b^hi^Ly6G^hi^) were examined for EdU incorporation and Ki-67 expression. (b - d) WT and Il4rα^-/-^ mice were inoculated with Nb for 3 days. (b) lung neutrophil EdU incorporation (3hrs after EdU injection) and Ki-67 expression, and p-STAT6 (c) and IL-13 (d) intracellular staining were assessed. Individual flow cytometry histograms and mean fluorescence intensity (MFI) are shown (a-d) for 3 mice/treatment group and are representative of 2 independent experiments. Flourescence Minus One (FMO) controls were used for background signal including gating of positive populations. Each symbol represents an individual mouse and horizontal lines indicate the mean. *p<0.05, **p<0.01, ***p<0.001, ****p<0.0001 (Student T test (a); one way ANOVA (b - d)).

## IL-4R signaling is required for the proliferative neutrophil phenotype

IL-4R signaling is a major signaling pathway that drives type 2 immunity and orchestrates characteristic activation states and associated effector and regulatory functions in macrophages and lymphoid cells after helminth infection^29^. To assess whether neutrophil activation and proliferation in the context of a type 2 immune response was dependent on IL- 4R signaling, *Il4rα*^-/-^ and control Balb/c WT mice were inoculated with Nb. At day 3 after inoculation, EdU was administered i.p. and 3 hours later lung neutrophils were assessed for proliferation. As shown in Fig. 2b, increases in both EdU^+^ and Ki-67^+^ neutrophils were blocked in Nb-inoculated *Il4rα^-/-^* mice. A small population of neutrophils in untreated mice also were observed to incorporate EdU and stain positive for Ki-67 (Fig. 2b). Few studies have yet examined effects of IL-4R signaling on neutrophils. To directly address whether IL-4R signaling was occurring in this myeloid subset, WT mice were stained with an antibody that specifically recognizes signal transducer and activator of transcription 6 (STAT6) when phosphorylated on tyrosine 641. STAT6 is specifically phosphorylated downstream of IL-4R signaling, thereby serving as a marker for IL-4R signaling^30, 31^. Expression of phosphorylated STAT6 (pSTAT6) by neutrophils was assessed at day 3 after Nb inoculation and 3 hours after EdU i.p. administration, allowing simultaneous assessment by flow cytometry of IL-4R signaling and cell cycling. As shown in Fig. 2c, pSTAT6+ neutrophils were markedly increased in EdU^+^ compared to EdU^-^ neutrophils. Surprisingly, although the number of EdU+ neutrophils was quite small in naïve mice, they still showed significant increases in pSTAT6 relative to naïve EdU^-^ neutrophils (Fig 2c), suggesting the presence of IL-4R signaling in a small proliferating lung tissue resident neutrophil population in naïve mice. Previous studies have shown that lung neutrophils express IL-13 after Nb inoculation^8^, making IL-13 a useful marker for neutrophil activation after helminth infection. To examine whether EdU^+^ neutrophils differentially express IL-13, in the same experiment as in Fig. 2b, neutrophils were stained for IL-13. As shown in Fig. 2d, IL-13 expression was markedly increased in EdU^+^ lung neutrophils in naïve mice and Nb inoculated mice relative to EdU^-^ neutrophil populations.

## Proliferative lung neutrophil subset is primarily c-kit+

Proliferation and oxidative phosphorylation are characteristic of a more immature neutrophil phenotype^32^, which in the periphery is now well-described in aberrant pro-cancerous tissues in the tumor microenvironment^5, 17^. As c-kit is a characteristic marker for immature neutrophils^17^, lung neutrophils at day 2 after Nb inoculation and 3 hours after EdU injection were analyzed for c-kit cell surface protein using flow cytometric analysis. As shown in the gating strategy (suppl Fig. 3a), c-kit+ neutrophils comprised about 3-4% of lung neutrophils at day 2 after Nb inoculation, and this population included the majority of EdU^+^ cells (Fig. 3a).

**Fig. 3.**
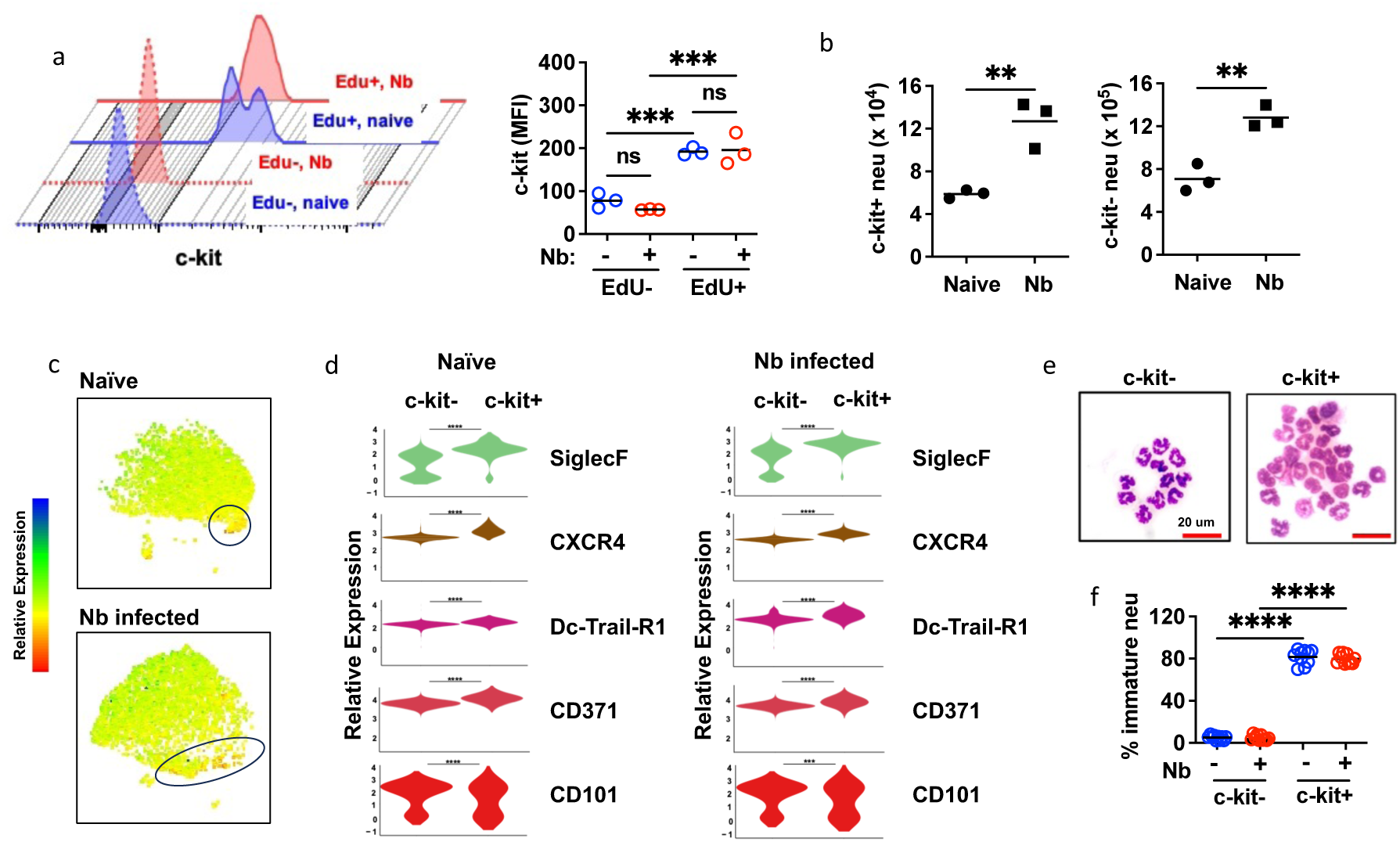
After Nb infection, proliferating lung neutrophils express c-kit and specific activation markers and are morphologically immature. (a-b) Mice were inoculated with Nb for 2 days with EdU injection (i.p.) for 24 hrs. (a) Representative c-kit expression histogram and mean florescence intensity (MFI) for 3 mice/trt group. (b) total number of c-kit^+^ and c-kit^-^ neutrophils in the lungs of naïve and Nb inoculated (2 days) mice. (c) Spectral flow cytometry of neutrophil activation markers, with UMAP of c-kit expression (c) and (d) violin plots of surface marker expression of c-kit^+^ and c-kit^-^ lung neutrophils from naïve and Nb infected (day 2) mice (5 mice/trt group). (e) Representative Wright-Giemsa stain of sorted c-kit- and c-kit+ neutrophils from the lung of naïve or Nb infected (day 2) mice. (f) Neutrophils are enumerated in each field with a total of 5 fields examined in each slide and present as percentage of immature neutrophils to total neutrophils. Each symbol represents an individual mouse (a, b), or technical replicates from pooled samples (e), horizontal lines indicate the mean, and **p<0.01, ***p<0.001, ****p<0.0001 (Student T test (b); one way ANOVA (a, f))

C-kit+ neutrophils also increased over 2-fold by day 2 after inoculation, as did c-kit^-^ neutrophils (Fig. 3b). To further characterize c-kit^+^ neutrophils, spectral flow cytometry was used to assess their expression of markers associated with neutrophil activation and maturation states in lungs of both naïve and Nb-inoculated mice (Fig. 3 c,d). SiglecF, expressed in neutrophils found in adenocarcinomas^33^ and non-small-cell lung cancer^34^, was upregulated in c-kit+ neutrophils, as was DC-Trail-R1 and CD101, which have recently been found to characterize neutrophils activated in the tme^5^. The chemokine, CXCR4, was also upregulated. Along with c-kit, CXCR4 is a ligand for CXCL12, which is required for neutrophil binding to endothelial cells in the lung^23^.

As such, these cell cycling c-kit+ lung neutrophils show upregulated expression of several neutrophil markers associated with the tumor microenvironment and neutrophil localization in the lung. Our previous studies have shown that at day 2 after Nb inoculation, many lung neutrophils exhibit a toroid or ring-shaped nucleus, in contrast to more mature multi-lobed nuclei^8^. Histological analysis of sort-purified c-kit^+^ and c-kit^-^ neutrophils demonstrated that c-kit^+^ lung neutrophils were enriched in toroid immature neutrophils in both naïve mice and at 2 days after Nb inoculation (Fig. 3 e,f).

## c-kit^+^ neutrophils express increased mRNA for cell cycling and type 2 responses

To globally assess gene expression profiles, electronically sorted c-kit^+^ and c-kit^-^ neutrophil populations from lungs of mice at 2 days after Nb inoculation were subjected to bulk RNAseq analyses (Fig. 4a). It should be noted that we could not obtain sufficient lung neutrophils from untreated mice for bulk RNAseq of c-kit^+^ cells; instead, transcriptional analysis of untreated neutrophil subsets were analyzed by single cell RNA seq (scRNAseq), as described later. As shown in Fig. 4b, PCA showed marked global differences in gene expression between c-kit^+^ and c-kit^-^ neutrophils. The previously described neutrophil maturation score^35^, based on expression of genes associated with neutrophil maturation and differentiation, was used to compare c-kit+ and c-kit- lung neutrophils at 2 days after Nb inoculation. As shown in Fig. 4c, c-kit^+^ lung neutrophils were significantly less mature than c-kit^-^ lung neutrophils. Using IPA, c-kit^+^ lung neutrophils after *Nb* inoculation had enriched predicted gene expression of canonical signaling pathways associated with proliferation, including cell cycle checkpoints, mitotic metaphase and anaphase, and S phase, increased mitochondrial activity (oxidative phosphorylation) and eukaryotic translation (Fig 4d). Expression of individual genes in these pathways relative to c-kit^+^ neutrophils are presented as a chord diagram in Fig 4e. Canonical immune response genes were also assessed with genes typically associated with type 2 responses such as Arg1, Chil3, and IL-13 upregulated while genes associated with type 1 responses were reduced (Fig. 4f). In contrast, relative to c-kit+ neutrophils, c-kit- neutrophils showed increases in expression of genes associated with degranulation, and also expression of genes associated with type 1 immunity, including IL-12 signaling pathways and reactive oxygen species (Fig. 4d,g).

**Fig. 4.**
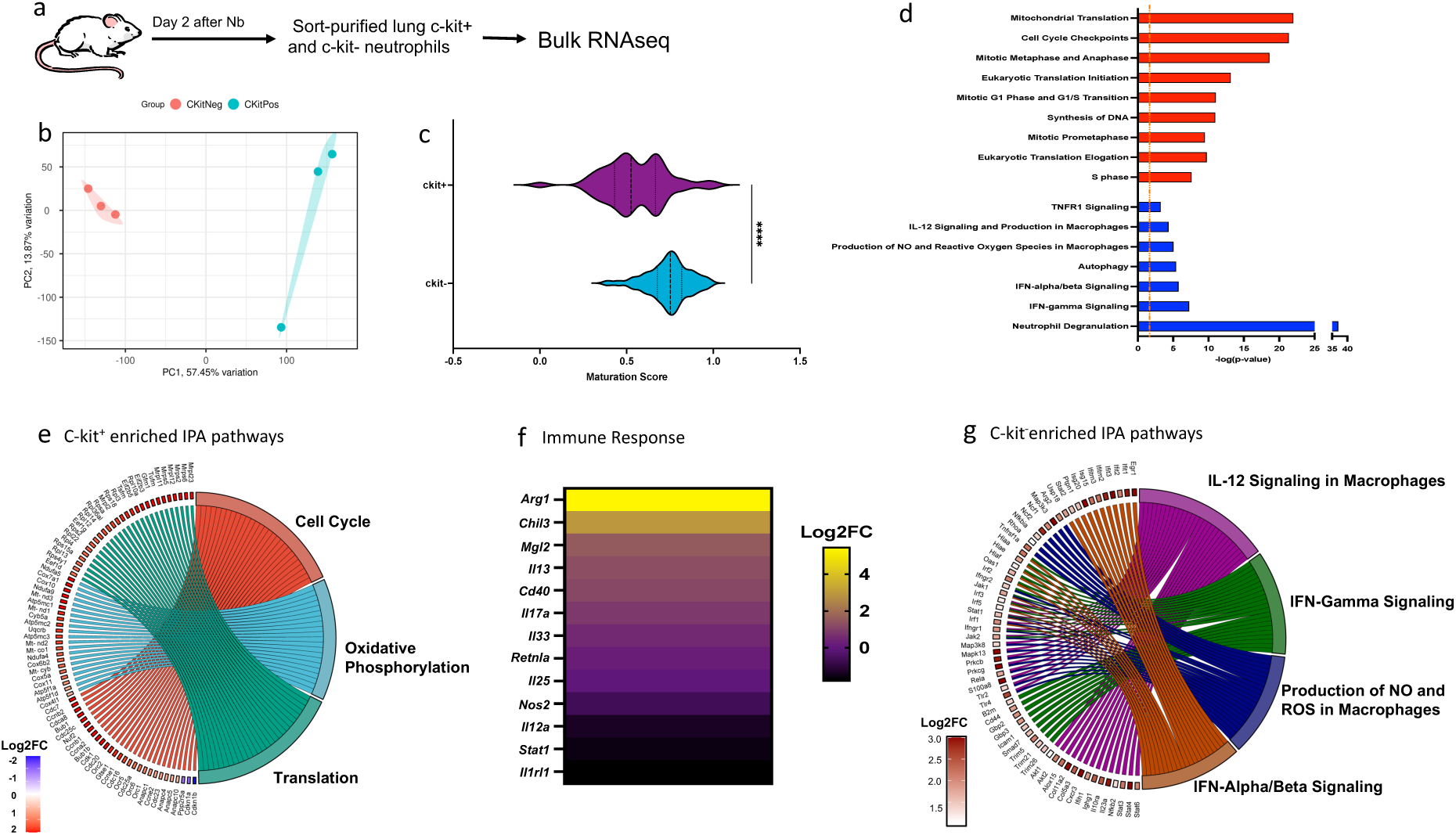
c-kit^+^ neutrophils show distinct upregulation of signaling pathways relative to c-kit^-^ neutrophils. (a) Mice (3/trt group) were inoculated with Nb for 2 days and lung c-kit^-^ and c-kit^+^ neutrophils were sorted and compared using bulk RNAseq. (b) Principal component analysis (PCA) of the normalized RNAseq data transcripts of c-kit^-^ and c-kit^+^ neutrophils. (c) Maturation score for c-kit^+^ and c-kit^-^ neutrophils. (d) Ingenuity Pathway Analysis (IPA) of differentially expressed canonical signaling pathways. Red bar indicates upregulated pathways in c-kit^+^ relative to c-kit^-^ neutrophils while blue color indicates suppressed pathways. Dashed orange line indicates log p<0.05. (e) Chord diagram shows differentially expressed genes (DEGs;c-kit^+^/c-kit^-^) related to upregulated IPA pathways. (f) Heatmap depicts expression of specific immune response genes (c- kit+/c-kit-). (g) Chord diagram shows DEGs upregulated in c-kit^-^ relative to c-kit^+^ neutrophils.

## Proliferating c-kit+ lung neutrophil population is initially localized in vascular endothelium

In the lung, in uninfected mice neutrophils occupy an endothelial cell niche binding to the capillary bed in part through Cxcl12-CxCR4 interactions^23^. Larvae begin to invade the lung tissue by 11 hours after infection^27^ and by 48 hours lungs are infiltrated with marked increases of neutrophils^10^. We used intravascular staining to localize c-kit+ proliferating neutrophils in the alveolar membrane of the lung and differentiate them from recirculating neutrophils in the pulmonary vasculature and neutrophils residing in the tissue capillary bed. At days 0 (untreated), 1 or 7 after Nb inoculation, mice were injected iv with FITC labelled anti-CD45 Ab and 3 minutes later mice were euthanized (Suppl Fig. 4a). Mice were not perfused to avoid artifactual redistribution of immune cells in the lung tissue microenvironments^36^. Cell suspensions from blood, BALF, and lung were counterstained for Ly6G and c-kit and analyzed by flow cytometry. As shown in Suppl Fig. 4b, in naïve mice, c-kit+ neutrophils were not detected in the blood, but were localized in the BAL and lung, while the c-kit+ iv neutrophils (CD45+) (Suppl Fig. 4c) were entirely restricted to the lung consistent with their being embedded in the capillary bed, and in many cases exposed to blood vessel circulation. At day 1 after Nb inoculation, c-kit+ neutrophils were already detected in the blood (Suppl Fig. 4d), and as expected, almost 100% were c-kit+ i.v. while c-kit+ neutrophils in the BAL did not express i.v.

CD45 Ab (Suppl Fig. 4e). Interestingly, in the lung about 40% of the neutrophils in lung tissue did not express i.v. CD45 Ab, indicating that these cells were not exposed to the circulation, likely a result of their migrating out of the endothelial niche as they enter the lung tissues in response to the invading parasites. Similar results were observed at day 7 after inoculation (Suppl Fig. 4 f,g). As neutrophil infiltration during Nb infection can cause lung hemorrhaging^10^, we tested for vascular leakage by injecting Evans Blue dye (EBD) as previously described^37, 38^. Although vascular leakage was not detected at day 1 or day 7, it was pronounced at day 3 (Suppl Fig. 4h). As such, intravascular staining was not performed at days 2-4.

To visualize potential migration of proliferating neutrophils from the capillary bed into the lung parenchyma, confocal imaging of lung tissue cryosections obtained at day 2 after Nb inoculation were compared to lung tissue cryosections from naïve mice. 24 hrs after EdU injection, proliferating (EdU^+^) neutrophils, which are c-kit^+^, in naïve mice were primarily localized and embedded within the endothelial vasculature (Suppl Fig. 4i,j; suppl. Fig. 5). After Nb infection, proliferating neutrophils were observed leaving the endothelial niche and entering the airways (Suppl Fig. 4k-n, Suppl. Fig. 5), consistent with the high percentage of c- kit+ noncirculating neutrophils (45%) detected in lung tissue by day 1 after Nb inoculation (Suppl Fig. 4e). At the site of parasite invasion of lung tissue, proliferating neutrophils were readily observed accumulating at the host/parasite interface (Suppl Fig. 4m).

## Single cell RNA sequencing (scRNAseq) reveals expansion of a distinct neutrophil subset after Nb inoculation

To extend these studies of neutrophil activation and heterogeneity in the lung, at day 2 after Nb inoculation, scRNAseq was performed on lung neutrophils from untreated and infected mice in the same scRNAseq assay. Individual analyses of four replicates for Nb-infected mice and untreated mice provided a robust interrogation of neutrophil heterogeneity. Unbiased analysis of the lung neutrophil dataset from both untreated and treated mice revealed 9 clusters (Fig. 5a), each of which expressed characteristic neutrophil-specific genes^12^(Suppl. Fig 6a). Differentially expressed genes in cluster 3, and cluster 8 evidenced a transcriptional program consistent with a more immature neutrophil maturation score (suppl. Fig. 6b). Also cluster 3, which expanded after Nb infection, showed upregulation of genes associated with cell cycling, survival and type 2 immunity and increased numbers of cells expressing these genes (Fig. 5b,c, Suppl Fig. 6c). Specifically, elevated mRNA expression of genes mediating cell cycling in cluster 3 included: *KI67*, *Cdc34*, *PolD4*, *Culb4*, *Yes 1*, and *Cdk6*. Type 2 genes upregulated in cluster 3 included *IL-4Ra*, *IL-13Ra1*, and *Arg1*. *IL-17Ra*, also upregulated, binds IL-17 which has recently been shown to support type 2 responses during Nb infection^39^. Interestingly, cells expressing elevated *Il4ra* were observed in all clusters, though still highest in cluster 3 (Fig. 5c). Average transcript density analysis for individual cells in naïve and Nb infected mice showed marked differences across clusters in both naïve and infected mice, indicating marked heterogeneity in neutrophil expression patterns across clusters (Suppl. Fig. 6d). Given the similarity in signaling pathways upregulated in cluster 3 and ckit+ neutrophils by bulk RNAseq, it appears likely that there is overlap between these two populations. It should be noted that c-kit mRNA was not detectable using scRNAseq (data not shown). *TRAILR1* and *CD101* were also upregulated in cluster 3, markers characteristic of cycling neutrophil subsets (T3) in the tumor microenvironment, which have been reprogrammed from recruited mature (T2) and immature (T1) neutrophils^5^. To reveal potential links between clusters in the lung microenvironment, we used a trajectory plot that orders cell differentiation based on RNA processing. RNA velocity indicated a pseudo time progression from clusters inclusive of 8, 0, and 4 to cluster 3 (Fig. 5d). This may indicate reprogramming in a heterogeneous population that may include tissue resident neutrophils and neutrophils recruited from the bone marrow, where helminth-induced type 2 inflammation drives the expansion of cluster 3 within the lung tissue microenvironment.

**Fig. 5.**
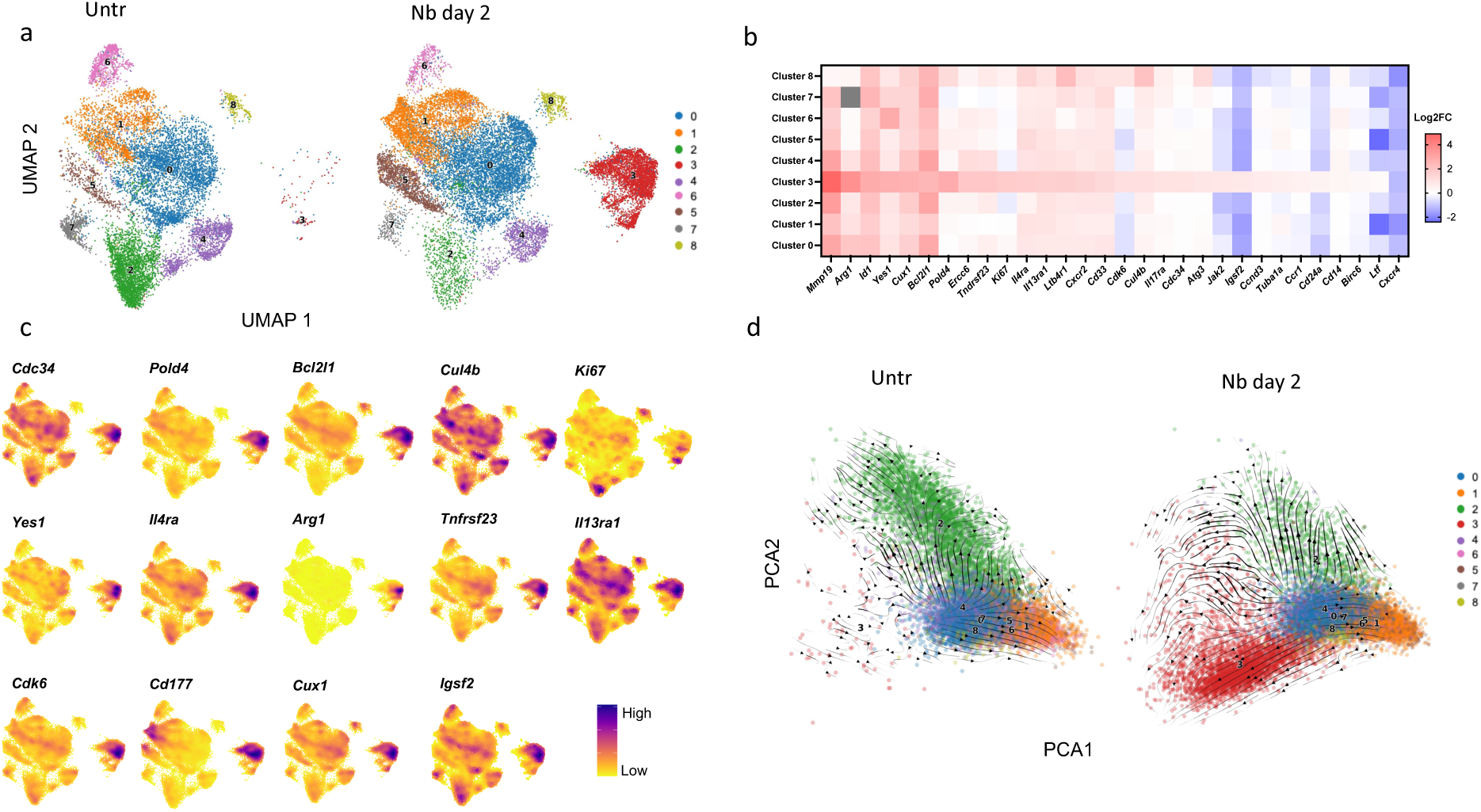
Lung neutrophil heterogeneity includes expansion of the proliferating cluster after *N. brasiliensis* (Nb) infection. Neutrophils were sorted from lung tissue cell suspensions collected from untreated or Nb inoculated (day 2) mice and transcriptionally analyzed by scRNAseq. (a) UMAP projection of neutrophils reveals nine clusters in naïve mice (left panel) and Nb infected mice (right panel). (b) Heatmap of markers related to cell cycling, immune responses, and wound repair in neutrophils of different clusters in Nb infected mice, as relative to neutrophils from naïve mice. (c) Integrated UMAP density plot representing density of cells expressing selected genes. (d) PCA plot of clusters shown in (a) with vectors indicating direction and magnitude of cellular transitions. Samples from individual mice (4/treatment group) were tracked by hash-tagging and individually analyzed.

## Type 2 immunity triggers a persistent proliferative neutrophil phenotype

Recent findings have shown adaptation of neutrophils to specific tissue environments including the lung^2^. Our findings now show that neutrophils exhibit broad plasticity of activation in the lung tissue microenvironment in response to pulmonary infections with distinct groups of pathogens. In direct experimental comparisons, our studies reveal that the lung neutrophil response to eukaryotic pathogens is distinct from responses to prokaryotes. Remarkably, following helminth infection, neutrophils undergo cell cycle progression and can persist for prolonged periods. This unique phenotype in lung neutrophils during infectious disease indicates functional differences that may contribute to host protection during type 2 immunity.

Although lung neutrophils from fungal infected mice did show pronounced mitochondrial activity relative to neutrophils from mice infected with bacteria, neutrophil cell death was similar in magnitude, and proliferation was not increased. Neutrophils play an important role as anti-fungal effector cells and also in promoting effector monocyte activation^40, 41^, functions that may require elevated metabolic activation. However, the potent type 2 immune response to helminth infection is quite distinct from the type 1 response triggered by microbial pathogens and the more mixed immune response resulting from fungal infection. Although Nb infection does initially promote emergency granulopoiesis, with pronounced IL-17 driven neutrophil infiltration of infected lung tissue, by 2-3 days after inoculation, the IL-17 response drops concomitant with an increase in IL-4 and IL-13^10^. Our findings indicate that the dominance of IL-4R signaling plays a major role in supporting the expansion of the proliferating neutrophil phenotype. Previous studies have shown that M2 macrophages proliferate during the type 2 response in peripheral tissues through IL-4R- dependent mechanisms^21^. Our studies now demonstrate that during the type 2 response neutrophils similarly require IL-4R signaling to proliferate in the lung tissue microenvironment. It indicates that activation of neutrophils can show parallel plasticity to macrophage activation during type 2 responses and demonstrates the impact of IL-4R signaling in driving activation of these distinct myeloid cell lineages.

The enrichment of proliferating cells and mitochondrial activity in the c-kit^+^ lung neutrophil subset after Nb inoculation is consistent with previous studies examining c-kit+ neutrophils in the bone marrow and in the tme^17^. Our studies now indicate that this population is also expanded in the lung during the type 2 immune response following Nb inoculation. C-kit^+^ neutrophils are generally considered a more immature population^17^ and our finding that this population was enriched in cells with toroidal nuclear morphology, characteristic of immature neutrophils rather than segmented nuclei^5^, suggests that it retains this immature state in the context of the lung microenvironment altered by helminth infection. The enhanced expression of type 2 markers, including Arg1 and IL-13, suggests that this population, though small relative to c-kit- neutrophils, may exhibit stronger functionality in the context of type 2 responses. Our observation that these proliferating neutrophils homed to the invading parasites in the lung tissue likely reflects their heightened role in mediating host protective responses. The expression of other markers by the c-kit+ neutrophils including dcTRAILR1, CD101, and SiglecF may also provide insights into their function. dcTRAILR1 and CD101 have been used as markers to identify more long-lived proliferative neutrophils in the tme^5^ while SiglecF can promote type 2 responses through release of cysteine leukotrienes^42^. Increased expression of Arg1 by neutrophils may indicate a role in mediating parasite killing, as Arg1 produced by macrophages has recently been shown to deplete Arginine availability to Nb L3, resulting in increased parasite mortality^43^. Our studies now show that helminth infection triggers a distinct neutrophil phenotype that undergoes proliferation, persists in the lung for prolonged intervals, and expresses markers similar to M2 macrophages. This remarkable plasticity in neutrophil activation likely helps to promote a host protective response tailored to the specific infectious pathogen.

Extramedullary neutrophils have not previously been shown to proliferate outside of the cancer tumor microenvironment (tme). In the tme, components of type 2 immunity are also observed including M2 macrophages and neutrophils that can contribute to the development of an immunosuppressive and pro-tumorigenic microenvironment^5, 44^. Their ability to promote resolution of type 1 inflammation and stimulate tissue repair processes, which promote disease tolerance in response to helminth infection, may be hijacked by tumors to promote their growth and survival. Indeed, in the scRNAseq analyses, cluster 3 expressed elevated CD101 and dcTRAILR1, markers proposed to characterize a recently identified T3 neutrophil population in the tme, which is derived from reprogrammed immature and mature neutrophils ^5^. Our trajectory analyses also suggest that the cluster 3 neutrophils may be derived from convergent reprogramming in both mature and immature neutrophils. Our observation that neutrophils proliferate during type 2 immunity, and also express characteristic M2 markers indicates mechanisms through which they may have additive effects to M2 macrophage functions in terms of mediating wound healing, anti-inflammatory, and tissue growth and repair functions in the TME and at the site of multicellular parasite invasion.

Resource availability

Lead contact:

Requests for further information should be directed to William Gause

## Materials availability

This study did not generate unique reagents.

## Data and code availability

All original data is available in the manuscript or supplementary information. This manuscript did generate original code. Any additional data needed to re-analzye the data in this manuscript will be provided by the lead contact upon request.

## Supporting information

Supplemental Figure with Legends

## Acknowledgements

This work was support by the National Institutes of Health Grants: RO1AI69770, R01AI131634, and R01 DK113790.

## Author contribution

Conceptualization: WCG, FC, AR, DP, MS; Funding Acquisition: WCG, AR, MS; Methodology: FC, SB, VE, LJ, KW, WW; Supervision: WCG, AR, DP; Writing original manuscript: WCG, FC; Writing review and editing: WCG, FC, AR; Bioinformatics: AL, AM, EJ, LJ, SB.

## Key Resources Table

| REAGENT OR RESOURCE | SOURCE | IDENTIFIER | COLOR |
| --- | --- | --- | --- |
| **Antibodies** |  |  |  |
| anti-CD45 (30-F11) | Biolegend | Cat. #: 103149, RRID: AB_2564590 | BV785 |
| anti-CD45 (30-F11) | Biolegend | Cat. #: 103108, RRID: AB_312972 | FITC |
| anti-F4/80 (T45-2342) | BD Biosciences | Cat. #: 749282, RRID: AB_2873657 | BUV805 |
| anti-CD64 (X54-5/7.1) | Biolegend | Cat. #: 139314, RRID: AB_2563903 | PE/Cyanine7 |
| anti-CD11c (HL3) | BD Biosciences | Cat. #: 612797, RRID: N/A | BUV737 |
| Anti-siglec-F (E50-2440) | BD Biosciences | Cat. #: 565526, RRID: AB_2739281 | PerCP-Cy5.5 |
| anti-CD11b (M1/70) | BD Biosciences | Cat. #: 565976, RRID: AB_2738276 | BUV395 |
| anti-CD11b (M1/70) | Biolegend | Cat. #: 101206, RRID: AB_312788 | FITC |
| anti-Ly6C (HK1.4) | Biolegend | Cat. #: 128024, RRID: AB_10643270 | AF700 |
| anti-Ly-6G (1A8) | Biolegend | Cat. #: 127636, RRID: AB_2563207 | AF594 |
| anti-Ly-6G (1A8) | BD Biosciences | Cat. #: 563979, RRID: AB_2738520 | BV711 |
| anti-c-kit (ACK2) | Biolegend | Cat. #: 135120, RRID: AB_2562040 | BV605 |
| anti-c-kit (ACK2) | BD Biosciences | Cat. #: 75269, RRID: N/A | BV605 |
| anti-CXCR4 (2B11/CXCR4) | BD Biosciences | Cat. #: 755964, RRID: N/A | RB780 |
| anti-CXCR2 (SA044G4) | Biolegend | Cat. #: 149313, RRID: AB_2734211 | APC/Cyanine7 |
| anti-DcTRAIL-R1 (mDcR1-3) | Biolegend | Cat. #: 133804, RRID: B_1626110 | PE |
| anti-CD101 (Moushi101) | eBioscience | Cat. #: 17-1011-82, RRID: AB_2815082 | APC |
| anti-CD115 (T38-320) | BD Biosciences | Cat. #: 751271, RRID: AB_2875286 | BUV615 |
| anti-CD206 (C068C2) | Biolegend | Cat. #: 141712, RRID: AB_10900240 | AF647 |
| anti-CD14 (rmC5-3) | BD Biosciences | Cat. #: 740125, RRID: AB_2739882 | BV510 |
| anti-CD371 (5D3/CD371) | BD Biosciences | Cat. #: 564795, RRID: AB_2738955 | BV421 |
| anti-CD3e (145-2C11) | BD Biosciences | Cat. #: 553061, RRID: AB_394595 | FITC |
| anti-CD19 (eBio1D3) | eBioscience | Cat. #: 11-0193-82, RRID: AB_657666 | FITC |
| anti-CD45.2 (104) | BD Biosciences | Cat. #: 562895, RRID: AB_2737873 | BV421 |
| anti-CD45.2 (104) | BD Biosciences | Cat. #: 564616, RRID: AB_2738867 | BUV395 |
| anti-CD45.1 (A20) | Biolegend | Cat. #: 110737, RRID: AB_2562565 | BV605 |
| anti-CD45.1 (A20) | Biolegend | Cat. #: 110715, RRID: AB_313504 | APC/Cyanine7 |
| anti-CD45.1 (A20) | Biolegend | Cat. #: 110731, RRID: AB_10896425 | BV421 |
| anti-CD31 (MEC13.3) | Biolegend | Cat. #: 102514, RRID: AB_2161031 | AF488 |
| anti-CD31 (390) | BD Biosciences | Cat. #: 740879, RRID: AB_2740529 | BV786 |
| anti-Ki-67 (16A8) | Biolegend | cat. #: 652406, RRIB: AB_2561929 | APC |
| anti-IL-13 (eBio 13A) | invitrogene | cat. #: 12-7133-82, RRIB: AB_763559 | PE |
| anti-STAT6 (J71-773.58.11) | BD Biosciences | cat. #: 558243, RRIB: N/A | AF488 |
| anti-STAT6 (J71-773.58.11) | BD Biosciences | cat. #: 558252, RRIB: AB_647233 | PE |
| anti-CD16/CD32 | Biolegend | Cat. #: 101302, RRID: AB_312800 | N/A |
| Zombie<sup>UV</sup> | Biolegend | Cat. #:423107, RRID: not available | amine reactive fluorescent dye |
| Evans blue dye | Sigma-Aldrich | cat#E2129, |  |
| Hi-Di formamide | Applied Biosystems | cat#43-113-20 |  |
| Click-iT EdU assay kit | ThermoFisher Scientific | cat#C10424 |  |
| 5-Ethynyl-2'-deoxyuridine (EdU) | TCI America, | cat# E1057 |  |

## Material and methods

### Mice

BALB/c ByJ (CD45.1), BALB/c (CD45.2), *Il4ra^−/−^* BALB/c mice and C57BL/6 mice were purchased from The Jackson Laboratory (Bar Harbor, ME), and were all bred and maintained in a specific pathogen-free, virus Ab-free facility at Rutgers New Jersey Medical School Comparative Medicine Resources. Healthy 8–12-week-old mice were selected for treatment groups from purchased or bred colonies, without using specific randomization methods or specific blinding methods. The studies have been reviewed and approved by the Institutional Animal Care and Use Committee at Rutgers-the State University of New Jersey. The experiments herein were conducted according to the principles set forth in the Guide for the Care and Use of Laboratory Animals, Institute of Animal Resources, National Research Council, Department of Health, Education and Welfare (US National Institutes of Health).

### Parasite culture, infection of mice, and 5-Ethynyl-2’-deoxyuridine (EdU) incorporation assay

*N. brasiliensis* (Nb) L3 were maintained in a petri dish culture containing charcoal and sphagnum moss. The larvae were isolated from cultures using a modified Baermann apparatus with 400U penicillin, 400 μg ml*^−^*^1^ streptomycin, and 400 μg ml*^−^*^1^ Neomycin (GIBCO, Rockville, MD) in sterile PBS, and then washed with sterile PBS three times.. During infections, mice were anesthetized with ketamine / xylazine and inoculated subcutaneously with a 100µl suspension of 500 Nb L3. *S. aureus* strain USA300 was administered intranasal with 2x10^7^ cfu. The fungi strain *A. fumigatus* CEA10 was administered by intratracheal inoculation with 3 x10^7^ live conidia per mouse.

For EdU incorporation in vivo, EdU (TCI America, Portland, OR, cat# E1057, 5 mg / ml, 100 ul in PBS) was administered to mice intraperitoneally (i.p.) 3 hours or 1 day before mouse sacrifice. EdU incorporation was assessed using Click-iT EdU flow cytometry cell proliferation assay kit following its protocol (ThermoFisher Scientific, Parsippany, NJ, cat#C10424).

### Flow cytometry, adoptive transfer, cell cytospin and Wright-Giemsa stain

Lung tissue was washed with stirring at room temperature for 10 min in Hank’s balanced salt solution (HBSS) with 1.3 mM EDTA (Invitrogen, Carlsbad, CA), then minced and treated at 37°C for 30 min with collagenase (1 mg / ml; Sigma-Aldrich, St. louis, MA, caty#381101) in RPMI1640 with 10% fetal calf serum (FCS), and then with 100 μg/ml of DNase for 10 min. Cells were lysed with ACK (Lonza, Walkersville, MD) to remove erythrocytes. Cells were blocked with Fc Block (CD16/32), directly stained with fluorochrome-conjugated antibodies against CD45, F4/80, CD64, CD11c, Siglec-F, CD11b, Ly6C, Ly6G, c-kit, CXCR4, CXCR2, Dc-Trail-R1, CD101, CD115, CD3, CD19, and Zombie^uv^,(see antibodies table) and analyzed by flow cytometry using either BD Biosciences lsrfortessa x-20 cell analyzer, or BD FACSymphony A5 high-parameter spectral cell analyzer (Franklin Lakes, NJ), and data analyzed by FlowJo (Ashland, OR). Cell sorting is conducted using BD FACSAria Fusion Flow Cytometer.

For adoptive transfer, both the donor CD45.2 mice and CD45.1 recipients were inoculated with Nb for 2 days. Neutrophils (Zombie^uv^-CD45^+^CD11b^hi^Ly6G^hi^) from the lungs of donor mice were electronically sort purified (>98%), 5 million cells were transferred i.t. into recipient mice. Ten days after cell transfer, the recipients received Edu (i.p.) as mentioned above for 24 hrs, sacrificed, and single cells were isolated from whole lung or collected from blood (400ul) by cardiac puncture and directly stained with fluorochrome-conjugated antibodies against CD45, CD11b, Ly6G and CD45.2. Cells were then fixed and permeabilized, using a kit from BD BioSciences (Franklin Lakes, NJ, cat# 554714). Half the cells were further stained intracellularly with a fluorochrome-conjugated antibody against Ki-67 or assayed for EdU incorporation as described above. All stained cells were analyzed by flow cytometry and gated on live, CD45^+^, CD11b^+^, Ly6G^+^ cells, then further gated on CD45.2 ^+^ neutrophils for EdU incorporation and ki-67 expression.

For cytospin analyses, neutrophils were sorted electrically from a pool of 6 mice from naïve, and 3 mice from Nb infection on day 2. Sorted cells (2 x10^5^) from lung tissue were suspended in 200 μl of 1x PBS with 2.5% FCS. The sorted cell suspensions were loaded into a Shandon Cytospin 4 (Thermo Electron Corporation, Waltham, MA), spun at 800- 1000 rpm for 5 min, and stained with Camco Quik Stain ® II, following its protocol (Cambridge Diagnostic products, Inc, Fort Lauderdale, FL, cat#211). Cells were visualized under a Nikon Eclipse 50i routine brightfield microscope equipped with the Q imaging Micropublisher 6 five MP USB 3.0 digital camera under 60x magnification. Immature neutrophils were defined as mono-nuclei, donut nuclei, and U shape nuclei; mature neutrophils were defined as nuclei having more than 3 segments (3 to 5 segment)^45, 46^.

### Lung neutrophil apoptosis/necrosis assay

Mice were infected with Nb, AF, or SA for 3 days, sacrificed, and lung tissue single cell suspensions were stained with antibodies to specifically detect neutrophils (CD45^+^, CD11b^+^, Ly6G^+^). Cells were then washed with cell staining buffer, further stained with Annexin V / propidium iodide kit (BD, San Diego, CA, cat#556547), following the manufacture’s protocol, and analyzed by flow cytometry.

### Intravascular staining of neutrophils and Evans blue dye staining to assess vascular leakage

Naïve or Nb infected mice under ketamine / xylazine anesthesia were injected intravenously (i.v.) with 3 ug of FITC - conjugated anti-CD45 antibody or its isotype control for 3 min. 300 ul of blood was withdrawn by cardiac puncture, followed by collection of BAL and lung tissue. For Ab staining, lung tissue single cell suspensions were prepared, RBC lysed and samples were Ab stained and analyzed via flow cytometry as described above. For the Evans blue dye (EBD) in vivo assay to test blood vessel leakage, naïve or Nb infected mice under ketamine / xylazine anesthesia were injected intravenously (i.v.) with 0.5% of EBD (Sigma-Aldrich, St. louis, MA, cat#E2129,) for 3 min. 300 ul of blood was withdraw by cardiac puncture, serum collected and lung tissue was transferred to a 2 ml size tube and dried at 57^0^C for 24h to 48h. The tissue was then weighed, minced, and 500 ul of formamide (Applied Biosystems, Foster City, CA, cat#43-113-20) was added to the tube. All tubes was transferred to a 55^0^C heat block and incubated for 24-48 hrs to extract EBD from tissue. Standard serial dilution of EBD was measured at absorbance at 610 nm, to calculate ng EBD extravasated per mg tissue, and further normalized as fold changes vs naïve mice, as previously described^37^.

### Immunohistological staining

Lungs were perfused with 200μl of PBS and Tissue-Tek O.C.T. compound (Sakura, Torrance, CA, cat#4583), excised and frozen in chilled acetone. 20 μm tissue sections were obtained using a HM505E cryostat (Microm International GmbH, Waldorf, Germany), 2% of rat serum for 1 hr, then stained with antibodies specific to CD31-FITC, Ly6G-AF594 for 1 hr after three washes with cell staining buffer. Tissues were fixed for 20 min. followed by permeabilizing for 20 min using the Click-iT EdU flow cytometry cell proliferation assay kit (ThermoFisher Scientific, cat#C10424), followed by the overnight Click-iT reaction at 4° C, and finally 90 min. Hoechst33342 (Invitrogen, Carlsbad, CA, cat#2064462) nucleus staining at room temperature. Coverslips were applied to the slides using ProLong Gold Antifade (Invitrogen, Waltham, MA, cat#2390722). Confocal images were taken using the Leica Stellaris 8 Stimulated Emission Depletion Super-Resolution microscopy system. Fluorescent channels were photographed separately, and images merged. Exposure times and fluorescence intensities were normalized to appropriate control images. Imaris interactive visualization and analysis software for 3D/4D microscopy images were used for image rendering and smart object detection.

### Bulk RNAseq and scRNAseq

For Bulk RNAseq, neutrophils from the lungs of naïve, Nb-, SA-, and AF-infected mice (2 days post inoculation) were sort-purified by gating on live, CD45^+^,CD11b^+^, Ly6G^+^ cells. The purity of all cell populations was 98% or greater. RNA was extracted using the RNeasy Plus Micro Kit (catalog no. 74034; QIAGEN). Illumina-compatible libraries were generated using the NEBNext Ultra II DNA Library Prep Kit for Illumina (catalog no. E7645S; New England BioLabs) and sequenced using an Illumina NovaSeq 6000 system in a paired-end 2x50-base pair (bp) reads configuration using the NJMS Genomics Center core facility. Bullk RNA-seq analysis was performed in accordance with the nf-core RNA-seq guidelines v.1.4.2. Briefly, the output reads were aligned to the GRCm38 (mm10) genome using STAR, followed by gene count generation using feature Counts and StringTie. Read counts were normalized and compared between groups for differential gene expression using DESeq2 with significance cutoff at false discovery rate-adjusted p<0.05. Determination of functional pathways was performed using Ingenuity Pathway Analysis (IPA) on differentially expressed genes.

For scRNAseq, lung tissue of naïve and Nb inoculated mice was collected and single cell suspensions prepared and sorted for neutrophils as described above. With Nb inoculated mice, samples from four mice were individually hashtag labelled (Totseq B0301 -B0308, Biolegend) and then pooled for sequencing. For untreated mice, due to low numbers of lung neutrophils, 4 samples (3 mice/sample) were individually hash tag labelled and then pooled for sequencing as described above. Hashtagged and pooled samples were processed for single cell sequencing on 10X Genomics platform using Chromium GEM-X Single Cell 3’ Reagent Kits v4 according to manufacturer’s protocol. Briefly, 29,000 flow sorted cells were loaded onto GEM-X chip for a targeted recovery of 20,000 cells followed by GEM generation and barcoding on Chromium iX instrument. After GEM-RT incubation, the samples were cleaned up and cDNA amplified. The amplified cDNA was purified using SPRI beads and separated into two fractions for preparing 3’ gene expression libraries and 3’ cell surface protein libraries (hashtag libraries). After library preparation and bead purification, each library was diluted to 4nM concentration. Gene expression libraries (GEX) were then pooled at 4nM concentration. Similarly, cell surface protein libraries (CSP) were also pooled at 4nM concentration. Both library pools were then diluted individually to 2nM and mixed at a ratio of 5:1 (GEX:CSP). The final library mix was sequenced on NovaSeq X Plus instrument at a final loading concentration of 170pM with 1% PhiX added according to Illumina Sequencing guide for 1.5B, 100 cycle kit. Paired end sequencing with run configuration Read1:28bp, Read2:90bp, i7 index:10bp and i5 index:10bp generated approximately 1 billion clusters per sample. Raw sequencing reads were barcode deconvoluted and aligned to the reference genome (mm10) via Cellranger (v 8.0.0). All subsequent processing was performed using the Seurat package (v 4.3.0) within R (v 4.2.2). Low quality cells (cells with percentage of reads of mitochondrial origin

>10%, with percentage of reads of ribosomal origin >45%, with percentage of reads of hemoglobin origin >20%, with <100 and >4000 feature counts, with <1000 counts of RNA) were filtered from the dataset, and read counts were normalized using the scTransform method. Samples were integrated with the Seurat integrate function, and clustered via UMAP according to nearest neighbors.

RNA velocity is a computational technique that leverages the ratio of spliced and unspliced RNA molecules to infer dynamic cellular state transitions from single-cell RNA sequencing data. In this study, RNA velocity analysis began with the generation of loom files using *velocyto* (v0.17.17) for each sample. The raw BAM files and a reference annotation file (refdata-gex-mm10-2020-A.gtf) were used as inputs to extract spliced, unspliced, and ambiguous transcript information. The splicing information from all samples was subsequently merged, and *scVelo* (v0.3.2) was employed to construct the RNA velocity graph. Transition probabilities across cells were estimated using cosine correlation between potential cell-to-cell transitions and velocity vectors. The calculated velocities were projected onto a lower-dimensional space using principal component analysis (PCA), with the resulting vectors indicating the direction and magnitude of cellular transitions. Finally, velocity pseudotime was computed to determine the progression of cells along inferred trajectories, and pseudotime was overlaid onto UMAP embeddings to visualize the temporal ordering of cells within the trajectory.

### Seahorse Assay for metabolic activity

Neutrophils from the lung of a pool of 3 mice were sort-purified as described above. Sorted neutrophils (∼75,000 cells/well) from each group were seeded into CellTak (Corning, New York, NY, cat# 354240) coated XFp plates in XF RPMI complete assay media and incubated at 37°C without CO2 for 45 minutes. The assay was run on a Seahorse XFp Extracellular Flux Analyzer (North Nillerica, MA) with each group of cells run in duplicate. After 20 cycles of basal measurement, Rotenone and Antimycin A (Seahorse XF Cell Mito Stress Kit), inhibitors of mitochondrial complex I and III, respectively, were injected into each well for a final concentration of 0.5 uM/well. Oxidative phosphorylation was measured by the oxygen consumption rate (OCAR; pmol/min) while the extracellular acidification rate (ECAR; mpH/min) was assessed by the accumulation of lactate, other metabolic acids and release of protons into the extracellular medium.

### Statistical analysis

Data were analyzed using the statistical software program Prism 8 (GraphPad Software, Inc., La Jolla, CA) and are reported as means (± SEM). Differences between two groups were assessed by student’s T-test, differences among multiple groups were assessed by one way ANOVA and individual comparisons were analyzed using Holm-Sidak test. Differences of p < 0.05 were considered statistically significant.

