## Supplemental Figure with Legends for "Helminth infection favors reprogramming and proliferation of lung neutrophils"

### Supplementary Figures with Titles and Legends

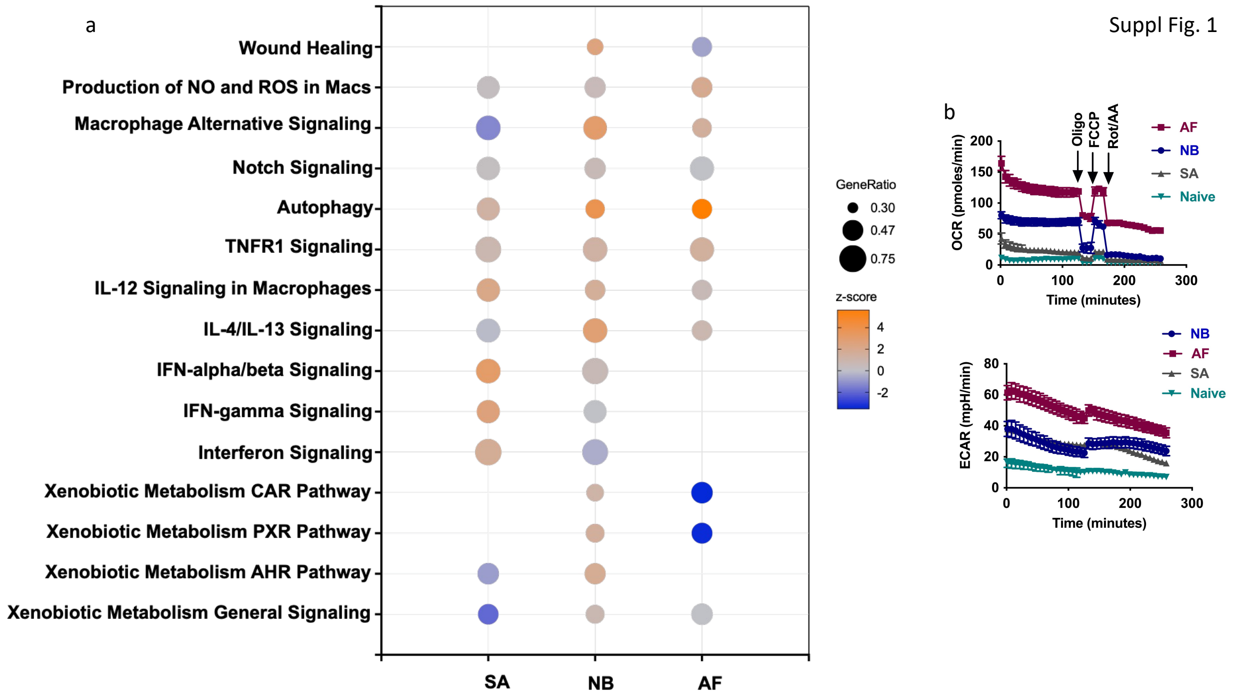


**Suppl. fig. 1. Lung neutrophil activation varies in response to infection with different types of respiratory pathogens**
Mice were inoculated with *N. brasiliensis* (Nb), *A. fumigatus* (Af), or *S. aureus* (SA) for 2 days. Lung neutrophils were sorted-purified for RNA-seq transcriptional analysis and compared to naïve neutrophils, with 3 mice/treatment group. (Left) Bubble plot of IPA pathway enrichment analysis of DEGs. "GeneRatio" indicates the ratio of enriched DEGs to total number of genes within the pathway. Z-score indicates activation status of the pathway. Orange signifies activation of the pathway, while blue shows inhibition. (b) Neutrophils were sort-purified and real-time metabolic activity assessed by Seahorse analysis. The oxygen consumption rate (OCR) and extracellular acidification rate (ECAR) were measured and inhibitors including Oligo, FCCP and Rotenone (Rot) and Antimycin A (AA) were added to block mitochondrial activity. Data shown were representative of two independent experiments.

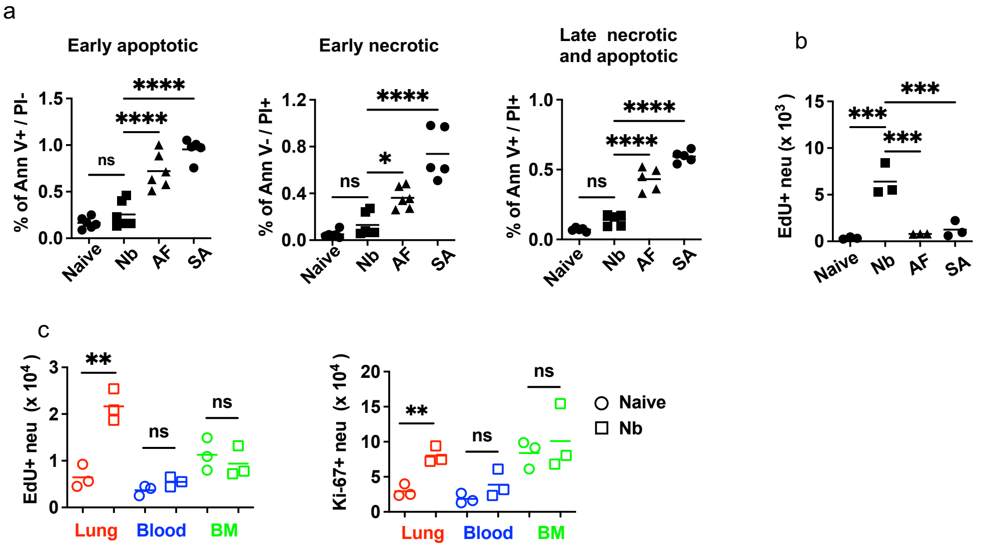


**Suppl. fig. 2. Neutrophils from helminth-infected mice exhibit reduced cell death and increased cell cycling**

Mice were infected with either *N. brasiliensis* (Nb), *A. fumigatus* (Af), or *S. aureus* (SA) for 3 days. (a) Neutrophils in lung cell suspensions were then stained for early apoptotic neutrophils (annexin ($V^+$/ $PI^-$), early necrotic neutrophils (annexin $V^-$/ $PI^+$), and late necrotic/apoptotic neutrophils (annexin $V^+$/ $PI^+$). (b) lung neutrophil EdU incorporation (3 hrs after administration) presented as total $EdU^+$ neutrophils. (c) Mice were infected with Nb for 3 days, and with EdU 3 hours before sacrifice. Neutrophils from the lung, blood, and bone marrow were analyzed for EdU incorporation or Ki-67 expression. Each symbol represents individual mice. All results are representative of two independent experiments. **p<0.01, ***p<0.001, ****p<0.0001 (one way ANOVA).

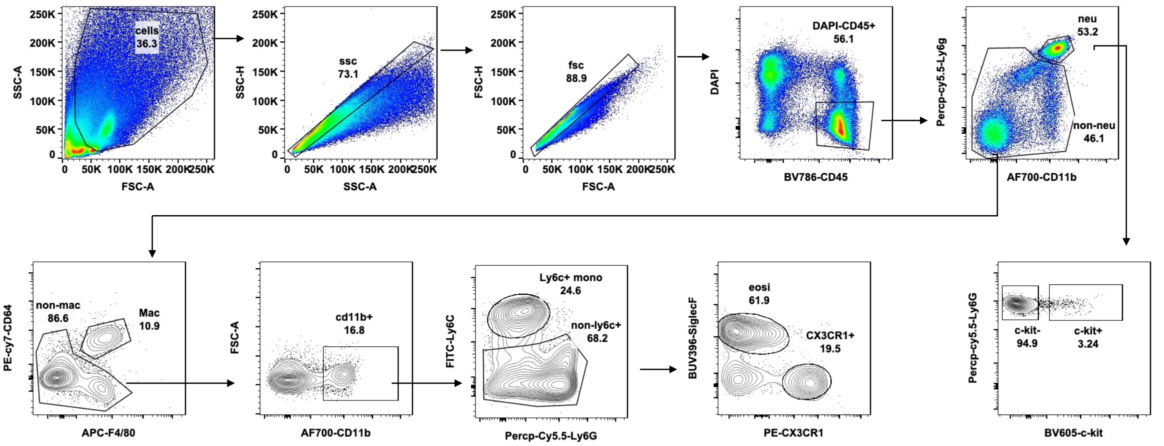


**Suppl. fig. 3. Gating strategy for flow cytometric analysis of lung neutrophil cell populations**

A representative flow cytometry gating strategy using lung samples from mice 2 days after Nb inoculation. Lung cell suspensions are gated on live single CD45$^+$ cells, which are then characterized as: Neutrophils: (CD11b$^{hi}$ Ly-6G$^{hi}$); c-kit$^+$ neutrophils: (CD11b$^{hi}$ Ly-6G$^{hi}$ c-kit$^+$). macrophages: (CD64$^+$F4/80$^+$, previous gating excludes neutrophils). Ly6C$^+$ monocytes: (CD11b$^+$Ly6C$^+$Ly6G$^-$, previous gating excludes neutrophils and macrophages). Eosinophils: (CD11b$^+$SiglecF$^+$CX3CR1$^-$, previous gating excludes neutrophils, macrophages, and Ly6C$^+$ monocytes). CX3CR1$^+$ monocytes: (CD11b$^+$CX3CR1$^+$Ly6C$^-$SiglecF$^-$).

Suppl. Fig 4

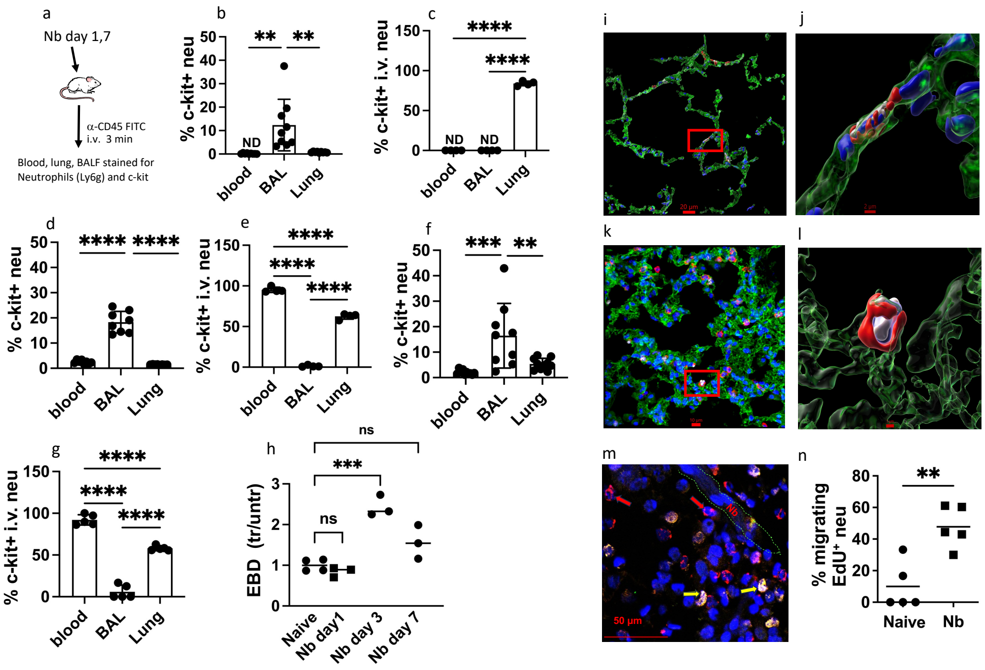


**Suppl fig. 4. Proliferating c-kit$^{+}$ lung neutrophils exit the lung vascular endothelium after *N. brasiliensis* infection.**

(a) Mice were inoculated with *N. brasiliensis* (Nb) for 1 or 7 days, injected intravenously (i.v.) with 3 ug of FITC-conjugated anti-CD45 antibody or its isotype control for 3 min., followed by BAL, blood, and lung tissue collection for flow cytometry. (b-c) untreated, % of neutrophils that are c-kit$^{+}$ (b), % of neutrophils that are c-kit$^{+}$ i.v. (c). (d-e) day 1, % of neutrophils that are c-kit$^{+}$ (d), % of neutrophils that c-kit+ i.v.(e). (f-g) day 7, % of neutrophils that are c-kit$^{+}$ (f), % of neutrophils that are c-kit$^{+}$ i.v. (g). (h) Mice were inoculated with Nb for 1,3, or 7 days, injected intravenously (i.v.) with 0.5% of Evan's blue dye (EBD) for 3 min. and concentration of the dye in lung tissue was measured and normalized as fold change over untreated mice. (i-m) Mice were inoculated with Nb for 3 days. Edu injection (i.p.) for 24 hrs. Lung tissue cryosections were stained with anti-CD31-FITC (green), anti-Ly6G Alexa Fluor 594 (red), EdU (white), and Hoechst 33342 nuclear counter stain (blue). Naïve (i, j) or Nb infected (k, l, m). Scale bar (20 μm in i, 2 μm in j, 10 μm in k, 1 μm in l, 50 μm in n). (n) % neutrophils migrating from endothelium, EdU+ neutrophils relative to total EdU+ neutrophils with each symbol representing an individual mouse, horizontal lines indicate the mean, and **p<0.01, ***p<0.001, ****p<0.0001 (one way ANOVA (b - h), or student t test (m)), ns: no significance.

Suppl Fig. 5

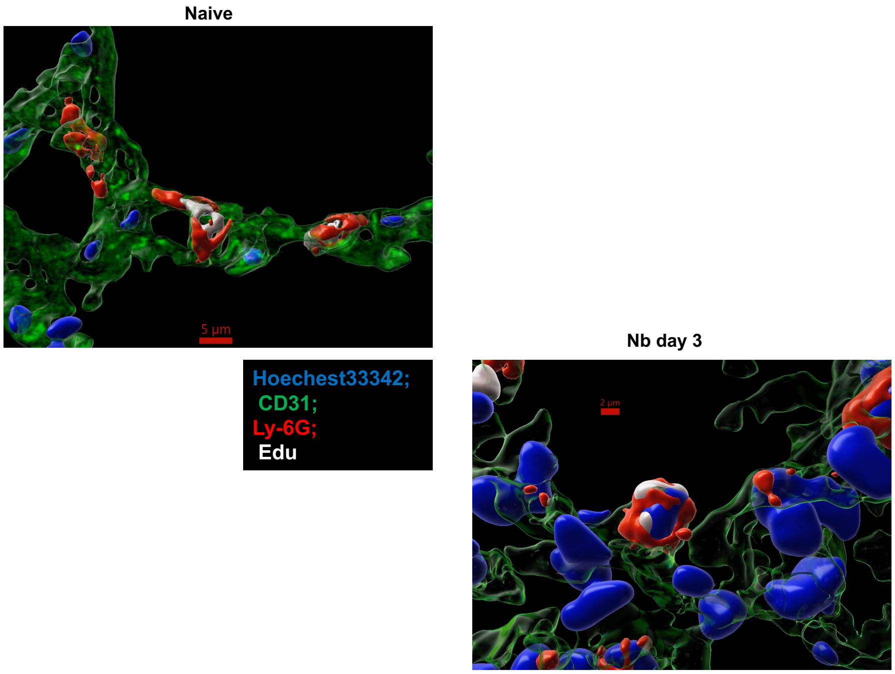


**Suppl. fig. 5. Proliferating lung neutrophil population is initially localized in vascular endothelium. (Actual videos are attached as MP4 files)**
Naïve mice and mice inoculated with Nb for 3 days were examined for proliferation using Edu incorporation.  Lung tissue cryosections were stained with anti-CD31-FITC (green), anti-Ly6G Alexa Fluor 594 (red), EdU (white), and Hoechst 33342 nuclear counter stain (blue). Confocal images were taken using the Leica Stellaris 8 Stimulated Emission Depletion Super-Resolution microscopy system. Fluorescent channels were photographed separately, and images merged. Exposure times and fluorescence intensities were normalized to appropriate control images. Imaris interactive visualization and analysis software for 3D/4D microscopy images were used for image rendering and smart object detection. Scale bar (5 µm in naive, 2 µm in Nb day 3).

Suppl. Fig. 6

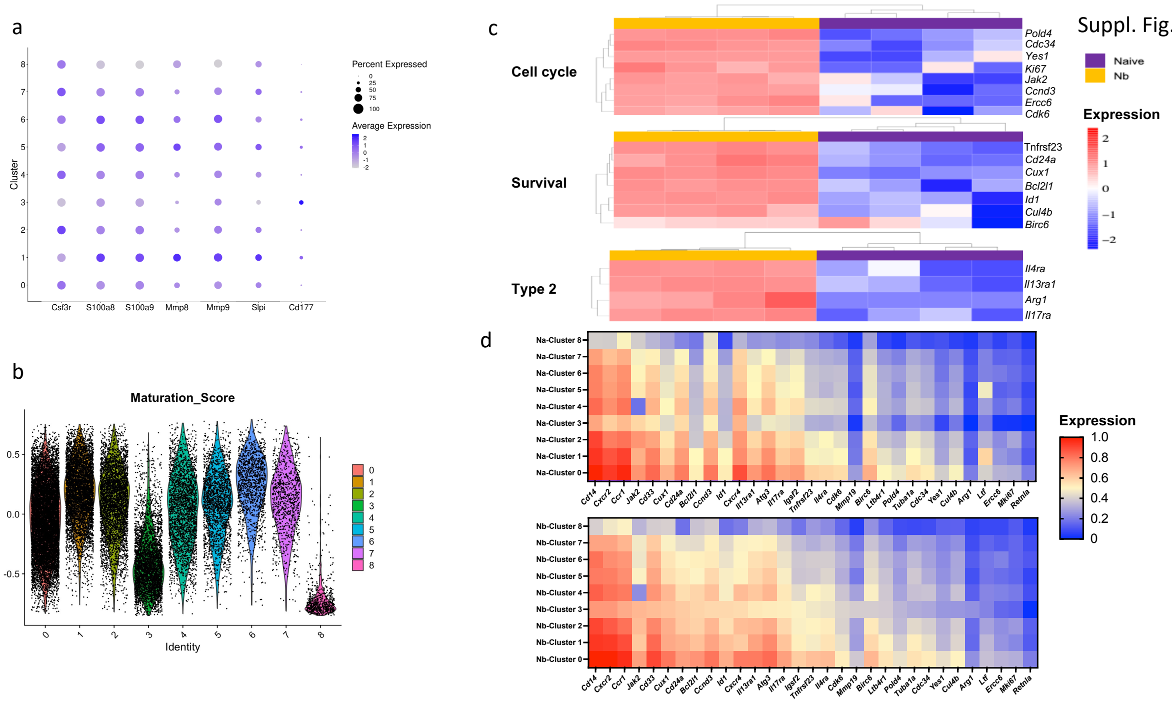


**Suppl. fig. 6. scRNAseq reveals expression of lung neutrophil genes in individual clusters, heterogeneity in maturation between clusters, and average expression of selected genes across clusters.**

Mice (4/trt group) were inoculated with *N. brasiliensis* (Nb) for 2 days and sort-purified neutrophils were transcriptionally analyzed using scRNAseq. (a) Dot plots show average expression of indicated genes characteristic of neutrophils, and the percentage of cells expressing the genes within each cluster. (b) Violin plots show module score of the gene maturation signature for each cluster identified in Figure 7a with scores closest to 0.5 being more mature. (c) Heatmap showing the average expression of genes in cluster 3 only for naïve and Nb-inoculated mice, with each column representing samples from individual mice. (d) Heatmaps representing average expression of selected genes (from Figure 7b) in naive and Nb infected mice.
